# Dimalis: A complete standalone pipeline to analyse prokaryotic cell growth from time-lapse imaging

**DOI:** 10.1101/2024.04.23.590675

**Authors:** Helena Todorov, Bouke Bentvelsen, Stefano Ugolini, Alan R. Pacheco, Anthony Convers, Tania Miguel Trabajo, Jan Roelof van der Meer

## Abstract

Real-time imaging of bacterial cell division, population growth and behaviour is essential for our understanding of microbial-catalyzed processes at the microscale. However, despite the relative ease by which high resolution imaging data can be acquired, the extraction of relevant cell features from images remains cumbersome. Here we present a versatile pipeline for automated extraction of bacterial cell features from standalone or time-resolved image series, with standardized data output for easy downstream processing. The input consist of phase-contrast images with or without additional fluorescence details, which are denoised to account for potential out-of-focus regions, and segmented to outline the morphologies of individual cells. Cells are then tracked over subsequent time frame images to provide genealogy or microcolony spatial information. We test the pipeline with eight different bacterial strains, cultured in microfluidics systems with or without nutrient flow, or on agarose miniature surfaces to follow microcolony growth. Examples of downstream processing in form of extraction of growth kinetic parameters or bistable cell differentiation are provided. The pipeline is wrapped in a Docker to facilitate installation, consistent processing and avoiding constant software updates.

## INTRODUCTION

Microorganisms play crucial roles in human, animal, and plant health, as well as in maintaining nutrient, water, and structural equilibria in environmental habitats. These roles ultimately rely on their spatial organisation, which influence the collective emergent properties of the multispecies communities they comprise. Understanding this behaviour at the level of single cells is crucial for determining how different environmental conditions and inter-organism interactions underly the growth of microbial populations.

A widely used way to gain insight into complex microbial behaviour is to cultivate single species or mixtures of microorganisms in synthetic growth media, and visualise the spatial and temporal development of individual cells via microscopy. Cells can hereto be embedded in microfluidic chambers representative of their natural habitat (Aleklett, et al., 2018; Dal Co, et al., 2020; Li, et al., 2014; Massalha, et al., 2017), or on nutrient surfaces (Hockenberry, et al., 2021; Reinhard and van der Meer, 2010; Seef, et al., 2021; Stylianidou, et al., 2016), which enable optimal focusing and tracking of individual cells. Taking successive images of the individual microbial cells at short time intervals (e.g., every few minutes) allows one to track their spatial positions, deduce cell division or demise, and observe or quantify specific individual physiological or phenotypic properties (e.g., by using fluorescent reporter gene fusions (Sulser, et al., 2022)). Microscopy time-lapse imaging has, for instance, provided unprecedented insight into the spatio-temporal development of microbiota in the plant rhizosphere (Massalha, et al., 2017), shown the interaction ranges of cross-feeding metabolites (Dal Co, et al., 2020), demonstrated the effect of substrate gradients on metabolic pathway expression (Tecon, et al., 2006), or illustrated the importance of subpopulation development in infection (Arnoldini, et al., 2014).

Whereas time-resolved microscopy allows to generate large data sets, in which thousands of individual microbial cells are imaged over tens to hundreds of time steps, the extraction of meaningful data is much more cumbersome. There is increasing need for automated image processing tools, which can identify individual cells on phase-contrast, differential interference contrast (DIC), or fluorescence images (*segmentation*), link cells across images taken at different time intervals, identify species from images or quantify specific markers that are representative for behavioural traits. The first cell segmentation and cell tracking tools appeared in the 1980s (Berns and Berns, 1982; Haralick and Shapiro, 1985; Kenong, et al., 1995), and established a foundation for the cell segmentation techniques that are still broadly used today (Carpenter, et al., 2006). More recently, however, the field has profited from the development of deep learning, with the appearance of highly accurate methods that can identify cells of variable size and shape with unprecedented levels of accuracy and in crowded environments (Cutler, et al., 2022; O’Connor, et al., 2022; Panigrahi, et al., 2021; Stringer, et al., 2021).

Among the plethora of image analysis methods that have been developed, users can choose between approaches that either require more manual user input and verification, or less input and more automatization. The more manual-oriented methods generally have a graphical user interface, and require the user to either set parameters or train models, which learn and improve cell-segmentation or -tracking until satisfactory results are achieved (Berg, et al., 2019; Carpenter, et al., 2006). More automated tools learn optimal segmentation on single focal strains, and permit user control on visual output or statistics from the identified cells (Meacock, et al., 2021; Padovani, et al., 2022; Stylianidou, et al., 2016). Other approaches attempt to limit user investment by training on large data sets of case images, optimising aspects of e.g., cell segmentation (Cutler, et al., 2022; Panigrahi, et al., 2021; Stringer, et al., 2021), denoising (Chen, et al., 2018; Izadi, et al., 2023; Liu, et al., 2018; Zhang, et al., 2017), or cell tracking (Hayashida and Bise, 2019; He, et al., 2017; Lugagne, et al., 2020). Tools such as Trackmate 7, aim to provide plug-ins to enable optimal transfer segmentation results to subsequent tracking tools (Ershov, et al., 2022). However, despite the enormous value of these tools, the field of procaryotic cell image analysis is, to our knowledge and experience, still lacking a complete, accurate, efficient and user-friendly pipeline, which enables all aspects of image analysis on large data sets: from raw image pre-processing, cell segmentation, cell tracking and data extraction, to representation of relevant cell features.

Here, we develop and test such a pipeline, which we named *Dimalis* (for *Di*gital *Ima*ge ana*l*y*sis*). Dimalis is provided as a *docker* image to facilitate installation, reproducible image analysis and data extraction with minimum user input. Basic computational skills are sufficient to install and run Dimalis. Dimalis is composed of individual published software tools, which we tested for optimal performance on different bacterial strains and conditions. The tools were embedded in an automated software environment, such that there is no need for successive intervention. After locating the raw image input, a single command is sufficient to start Dimalis and perform all subsequent tasks of image pre-processing, cell segmentation, cell feature extraction and cell tracking. *Dimalis_fluo* in addition enables extraction of single-cell fluorescence information (Fig. 1). Dimalis outputs can directly be used for common downstream processing software, such as network-visualisation, statistics or deep learning. We present here the pipeline with the individual components, and demonstrate the results of comparative tests on a variety of bacterial species imaged in time-lapse microscopy of surface-grown cells, microcolonies, subpopulations, and within microfluidic platforms. We are confident in the capabilities of Dimalis to substantially facilitate single cell analysis, and lowering the barrier of entry for researchers across biological disciplines.

**Figure 1:**
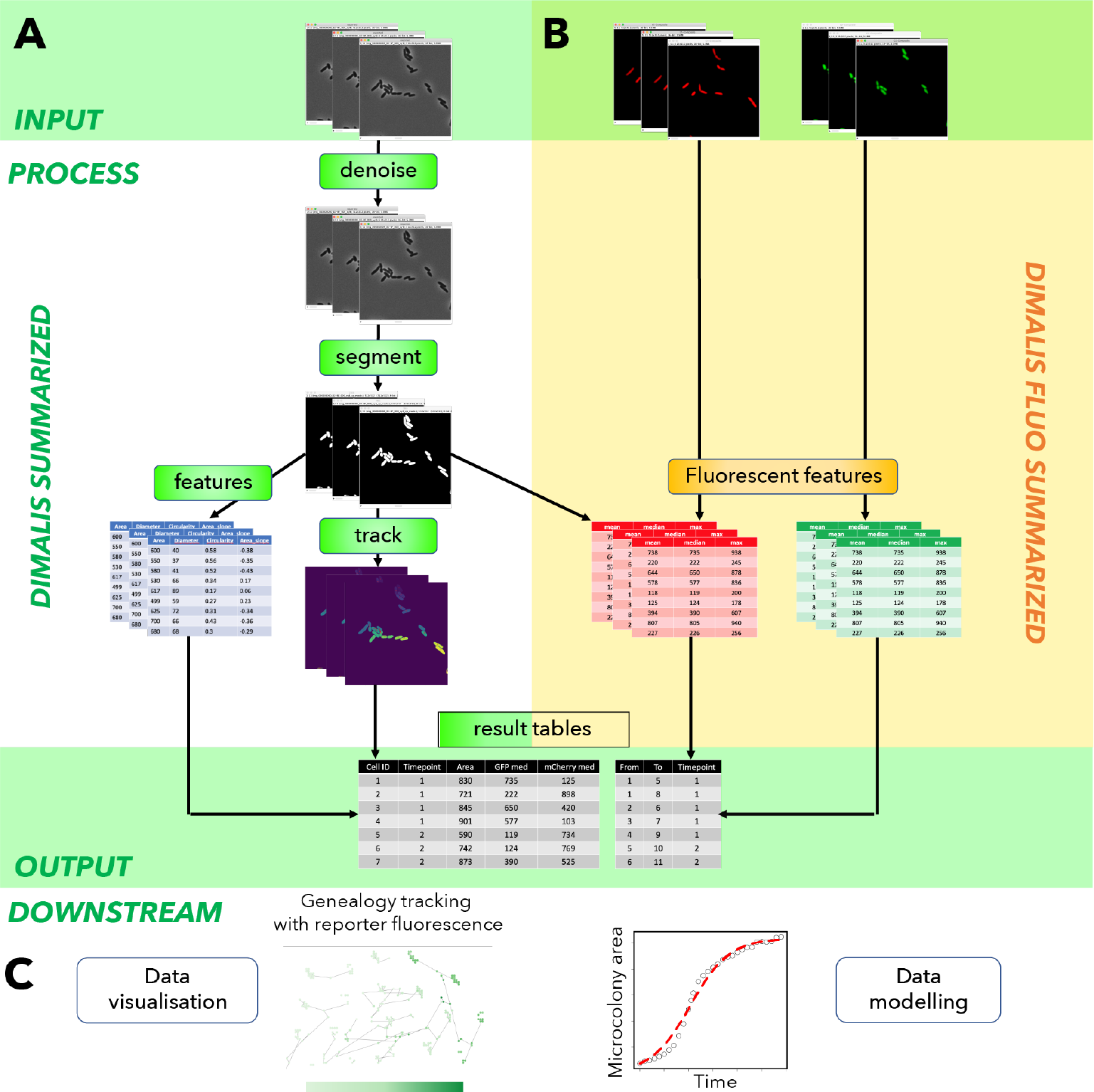
Overview of the Dimalis pipeline. A) Dimalis takes phase contrast or bright-field images as input, which are denoised and segmented, and from which cells are tracked and features extracted. The results are presented in forms of tables, which contain the features for all individual segmented cells all timepoints, and listing the corresponding cell and time tracks. B) Fluorescent cell image data as additional input for Dimalis. Fluorescence images will undergo a step of illumination correction, after which fluorescence data are quantified from every individual segmented cell in part A, and added to the summary tables. C) Summary tables can be directly used for downstream processing or visualization.

## MATERIALS AND METHODS

### Generation of images for automated tools comparison

The bacterial species *Pseudomonas putida* F1 (Zylstra and Gibson, 1989), and *Pseudomonas veronii* 1YdBTEX2 (Morales, et al., 2016) were used for ground truth comparisons. In addition, we cultured a diversity of soil bacterial isolates: *Lysobacter* sp., *Burkholderia* sp., *Curtobacterium* sp., *Mucilaginibacter* sp. and *Rahnella* sp. (all described in Ref (Čaušević, et al., 2022)). For imaging, cells were experimentally grown into monolayer microcolonies on miniaturized nutrient agarose surfaces as described previously (Reinhard and van der Meer, 2010). Cells were imaged on a Nikon ECLIPSE Ti Series inverted microscope coupled with a Hamamatsu C11440 22CU camera (Delavat, et al., 2016; Reinhard and van der Meer, 2010), and a Nikon CFI Plan Apo Lambda 100× Oil objective (1.45 numerical aperture) at 1000× magnification with intervals of 10 or 20 min depending on the strain, for a total duration of between 12 and 20 h. Cell images were saved as 16-bit .tif files. First, *P. putida, P. veronii, Lysobacter* and *Rahnella* images were manually analysed to generate ground-truth cell segmentation and tracking. For this, cell masks were manually traced on the phase-contrast images using the QuPath open-source software for bioimage analysis (Bankhead, et al., 2017). Cells were then manually tracked between consecutive time steps using the MaMut tracking Fiji plugin (Wolff, et al., 2018). After that, we automatically analyzed time-lapse images of *Burkholderia, Rahnella, Curtobacterium* and *Mucilaginibacter* with the Dimalis pipeline.

### Microfluidics systems and growth conditions

Microfluidic systems featuring microchambers for trapping and culturing bacteria were fabricated by photolithography of SU8 on silicon master moulds and subsequent soft lithography of PDMS, as previously described (Burmeister, et al., 2019). Microchambers of 1 μm height and 40×40 μm width allowed for growth of bacteria in monolayers, enabling imaging at single-cell resolution.

Leaf bacteria were sourced from the At-LSPHERE, a collection of 224 strains isolated from wild *Arabidopsis thaliana* leaves that represent a taxonomic and functional cross-section of the leaf microbiota (Bai, et al., 2015). Each strain was streaked from a glycerol stock stored at –80°C onto R2A agar (Sigma-Aldrich) supplemented with 0.5% (v/v) methanol. The plates were incubated at 22°C for 3 days, after which the cells were suspended (1 μl loop-full) in 2 ml R2A liquid medium (Sigma-Aldrich) supplemented with 0.5% (v/v) methanol in 20 ml culture tubes. Liquid cultures were incubated at 28°C angled at 45° with shaking at 220 rpm for 18 hours. Cultures were then washed three times via centrifugation at 6000 rcf for 3 minutes, vortexing, and resuspension in a 10 mM magnesium chloride solution.

A concentrated cell solution (10^8^ cells ml^−1^) was injected in the inlet holes of the microfluidic device and led to trapping of 1-10 cells in several microchambers. Subsequently, the device was connected to a continuous perfusion of a minimal medium containing 10 mM of xylan (Carl Roth) at 0.1 ml h^−1^ for 48 hours. The medium contained 2.4 g K_2_HPO_4_, 1.08 g NaH_2_PO_4_ · 2H_2_O, 1.62 g NH_4_Cl and 0.2 g MgSO_4_ · 7H_2_O per litre. The medium (1 L) was additionally supplemented with the following trace elements: 15 mg Na_2_EDTA · 2H_2_O, 3 mg FeSO_4_ · 7H_2_O, 4.5 mg ZnSO_4_ · 7H_2_O, 3 mg CoCl_2_ · 6H_2_O, 0.64 mg MnCl_2_, 1 mg H_3_BO_3_, 0.4 mg Na_2_MoO_4_ · 2H_2_O, 0.3 mg CuSO_4_ · 5H_2_O, and 3 mg CaCl_2_ · 2H_2_O. All media components were prepared with Milli-Q quality water (Millipore). Phase contrast images were taken at 40× magnification every 10 minutes by a Nikon Ti2 inverted microscope. Single cell elongation rates (*k*) were calculated, as previously described (Dal Co, et al., 2020), by fitting an exponential curve of the type *L*_*t*_ = *L*_0_2^*kt*^ to the length of the major axis of a cell over time. Tracks were filtered to keep only the ones tracked for longer than 4 timepoints. A tool for plotting growth parameters such as division events and elongation rates in space was coded by initially generating bins in x- and y-direction defined by the users (e.g., 15×15 μm). The tool then computes either the occurrences of division in each bin or the average elongation rates occurring in each bin, and returns a heatmap via the Seaborn Python library.

### Time-lapse imaging of transfer competent cells

To image the appearance of transfer competent cells in cell lineages of *P. putida* UWC1 -ICE*clc* (Sulser, et al., 2022), we used a double genetically tagged strain. First, we integrated a single copy mini-Tn*5*-integrated gene fusion between the *ahpC-*promoter and *egfp* to monitor induction of the alkyl hydroxyperoxidase as proxy for oxidative stress response (Delavat, et al., 2016). The resulting strain was then further tagged with a single-copy integrated P_inR_-*mcherry* reporter fusion to monitor activation of the Integrative and Conjugative Element (ICE) at single cell level. Bacterial cells were revived from –80 °C stocks and plated on nutrient agar plate containing 100 mg l^−1^ tetracycline and 25 mg l^−1^ kanamycin to select for the reporter constructs. A single colony was transferred and grown in 5 ml liquid LB media for 16 h at 30 °C, and then diluted 200 times in type 21C Minimal Media (MM) (Gerhardt, et al., 1981) supplemented with 10 mM sodium succinate (plus tetracycline and kanamycin), and again grown at 30 °C for 24 h. Cells were subsequently washed and diluted 100 times in MM without added carbon source. An aliquot of 6 μl of this suspension was spread on the surface of circular agarose patches (produced in MM and containing 0.5 mM 3-chlorobenzoate) enclosed in a black anodized POC (Perfusion Open and Closed) chamber (H. Saur Laborbedarf, Germany), which was mounted on a Nikon ECLIPSE Ti Series inverted microscope coupled with a Hamamatsu C11440 22CU camera (Delavat, et al., 2016; Reinhard and van der Meer, 2010). POC chambers were incubated at constant (22 °C) temperature and cells were imaged with a Nikon CFI Plan Apo Lambda 100× Oil objective every 30 min with an exposure time of 30 ms for phase contrast, 100 ms for GFP fluorescence, and 50 ms for mCherry fluorescence. Eight randomly selected positions per patch were followed during 43 h under the control of a script in MicroManager Studio (v1.4.23).

### Comparison of cell segmentation tools

We used two metrics to compare the results of automated cell segmentation tools to manual cell annotations. The F-score metric is the harmonic mean of precision and recall, computed as follows:

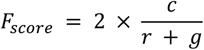

where *c* is the set of cells that were common between a tool’s results and manual annotations, *r* is the set of cells identified by the tool and *g* the set of cells that were manually identified. This metric thus provides information on both the precision 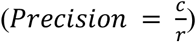 and recall 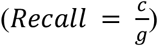. The Jaccard index measures pixel matching between the pixels identified as cells in manual annotations and in a tool’s results (Chenouard, et al., 2014). It is computed as follows:

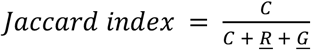

where *C* is the sum of pixels identified as cells in both manual annotations and the tool’s results, *<u>R</u>* is the sum of pixels identified as cells by the tool but not by the experts, and *<u>G</u>* is the sum of pixels identified as cells in the manual annotations but not in the tool’s results.

### Details of the Dimalis structure

Dimalis is wrapped in a *docker* structure. The pipeline contains the BM3D image denoising tool (Djurović, 2016; Sheng, et al., 2014) (version 4.0.1), the Omnipose cell segmentation algorithm (Cutler, et al., 2022) (version 0.7.1), the STrack cell tracking algorithm (Todorov, et al., 2023) (version 4), and home-made python scripts that allow to extract cell features using the regionprops *scikit-image* module from python.

### Testing different Omnipose versions

On a unique CPU compute node (2 x AMD Epyc2 7402) on the computational cluster from the University of Lausanne (with a 24 core processor), in a virtual environment, we always uninstalled all cellpose/omnipose previous versions from the *venv* before installing a new one. The cellpose/omnipose versions we tested are 0.7.0, 0.7.1, 0.7.2 and 2.2 (that we installed using the *pip install cellpose==0*.*7*.*0* command). All Cellpose/Omnipose versions were launched on the same phase contrast images, with the same set of parameters (using the pretrained model “*bact_omnitorch_0*” and a cell diameter of 29.5 every time).

### Data and code availability

The complete Docker version of Dimalis can be downloaded and installed from https://github.com/Helena-todd/Dimalis and https://github.com/Helena-todd/Dimalis_fluo.

## RESULTS

### Selection of automated segmentation and tracking tools

The extraction of precise cell information from time-resolved live single cell imaging relies on two crucial steps: segmentation and tracking. Cell outlines need to be correctly defined in each image, and the resulting cell masks need to be correctly linked between successive time points. First, we measured the precision of a variety of available segmentation and tracking tools on the same set of bacterial cell images, in comparison to a ground truth of manually segmented and tracked cells provided by experts in the field of microbiology. The images represented monocultures of four different bacterial strains, imaged over 16 time points. The manual segmentation and tracking resulted in two reference sets of cell masks and tracks, to which the results of the automated tools were then compared. The comparisons incorporated different metrics that allowed us to define, on a range of 0 to 1, the accuracy of the automation tools (Fig. 2).

**Figure 2:**
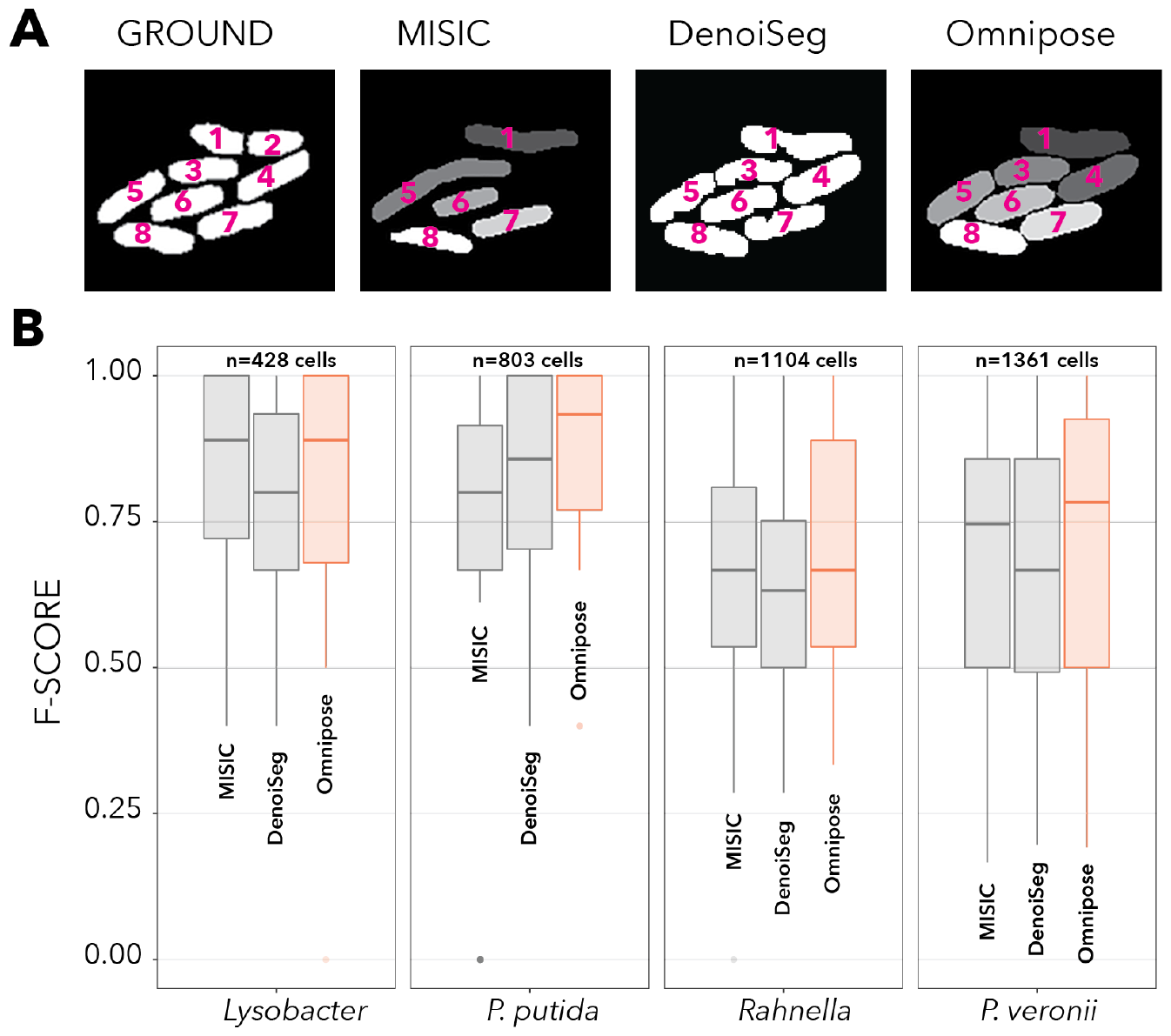
Comparison of automated cell segmentation tools to manual ground truth using the F-score. A) Manual segmentation of a microcolony of *Pseudomonas putida* cells (used as ground truth) in comparison to cell segmentations using MISIC, DenoiSeg or Omnipose. Segmented objects numbered according to ground truth. B) Overall accuracy (as F-score of the manual ground truth segmentation) for each of the three segmentation tools on n=5 replicate image sets of four bacterial strains (*Lysobacter* sp., *P. putida, Rahnella* sp. and *Pseudomonas veronii*). n, number of analyzed cells indicated on the top of each panel. An F-score of one corresponds to a perfect match between the automated and manual segmentations. Highlighted in orange is the tool with the highest averarge F-score for each bacterial strain tested. Boxplots show median, lower and upper quartiles, whereas whiskers indicate the 5^th^ and 95^th^ percentiles.

The F_score_ metric provides information on how accurately cells were retrieved by the automated cell segmentation tools, encompassing both the fraction of ‘true’ cells among all the cells returned by the tool (*precision*) and the fraction of true cells among the true number of cells (*recall*; see Materials and Methods). Cells of *Lysobacter* or *P. putida* were on average segmented more accurately than cells from *Rahnella* or *P. veronii*, which may be due to the latter two showing more crowded cell images than the other strains (Fig. 2B). Across all four bacterial strains, Omnipose had the highest F_score_.

Since the F_score_ only compares the fraction of true cells (Versari, et al., 2017), we used a second metric, the Jaccard index, to assess how well the cell mask contours were identified by each tool. Here we compared pixel matching between the manually drawn masks and the masks that were automatically identified (Fig. 3A). The Jaccard index (JI) typically returned lower values than the F_score_ for all tools and all bacterial strains except for *Rahnella* (Fig. 3B). For most bacterial strains, we observed comparatively lower accuracy of MISIC than for the other two automated tools, whereas Omnipose on average was again the best-performing tool. Consequently, we included Omnipose for cell segmentation in the Dimalis pipeline. For the cell tracking across time-lapse images we used our recently developed tracking tool STrack (Todorov, et al., 2023). The precision of STrack cell tracking in comparison to other tracking tools has been described in Ref (Todorov, et al., 2023).

**Figure 3:**
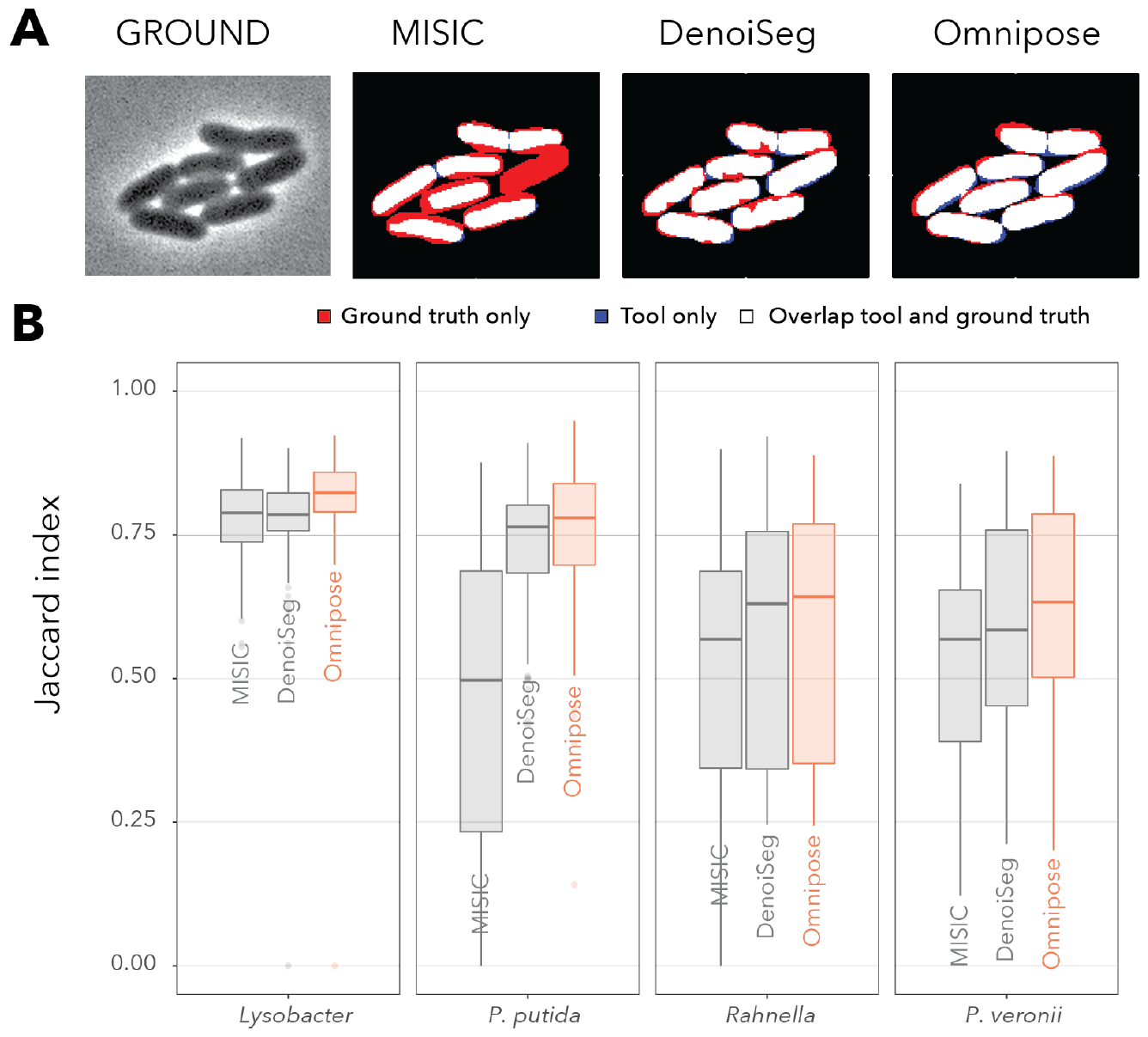
Comparison of MISIC, DenoiSeg and Omnipose using the Jaccard index. A) Manual segmentation of *P. putida* (ground truth) in comparison to automated cell segmentation using MISIC, DenoiSeg or Omnipose. Pixels identified as cells only in the ground truth are coloured in red in the others. Pixels identified as cells only by a tool are coloured in blue, whereas pixels identified as cells in both the ground truth and the tool are shown in white. B) Accuracy of cell outline matching for four different bacterial strains, shown as the Jaccard Index (proportion of matched pixels between the ground truth and the tool). A Jaccard index of one corresponds to a perfect overlap between ground truth and the tool’s segmentation results. Total number of between 428-1361 segmented cells (see Figure 2) on 5 replicate image positions with 16 time steps. Boxplots as in Figure 2.

### Image denoising

Time-lapse imaging of cells across long time spans of multiple hours often results in single images being slightly out of focus (despite automated focusing systems). An out-of-focus image can affect the cell segmentation results. We thus tested inclusion of an image denoising step to facilitate the detection of cell contours for those images where segmentation results would be sub-optimal (on the original images). Image denoising aims at reducing the noise and increasing the signal, which can help to correctly define objects. However, the sources of noise can be multiple, which significantly increases the complexity of the task at hand. As with all denoising techniques, there is also a trade-off between noise removal leading to signal-to-noise ratio increase, and noise removal leading to inevitable signal loss. Many denoising techniques exist and are available to the user, among which a variety is designed to correct specific sources of noise (Fan, et al., 2019). We tested here the BM3D denoising method that targets generic noise types (Chan, et al., 2017; Dabov, et al., 2007; Djurovic, 2016; Sheng, et al., 2014; Wang, et al., 2022). BM3D can reduce ‘salt and pepper’ noise, and will smooth flat pixel-value areas while enhancing cell contours, thus increasing the signal to noise ratio. The effect of this can be seen in Figure 4, with a cell example being slightly out of focus (Fig. 4A), and the denoised result (Fig. 4B). Cell segmentation was significantly poorer on the original image, retrieving only three cell masks and with significantly affected cell contours (Fig. 4C), whereas segmentation on the denoised image largely improved, retrieving four cells with more faithful cell masks (Fig. 4D).

**Figure 4:**
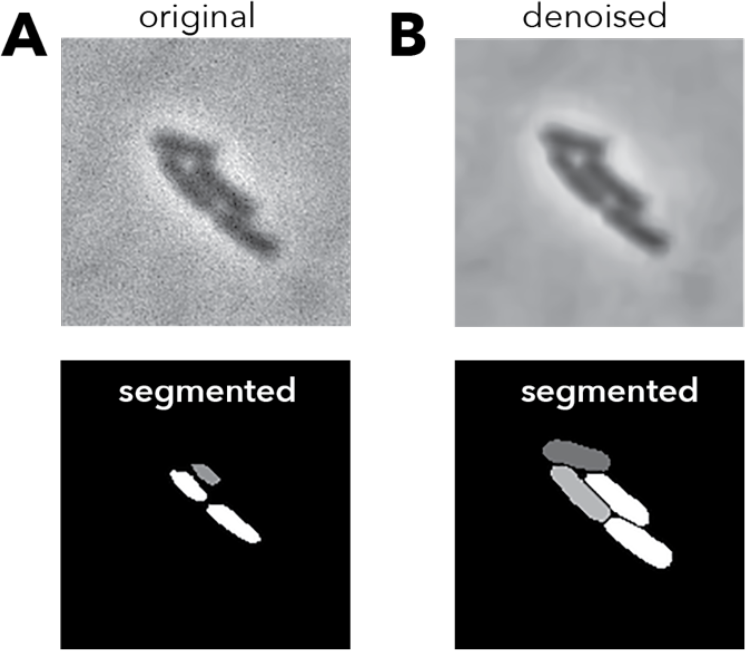
Denoising images improves cell segmentation. A) Original phase-contrast image of a microcolony of *Lysobacter* sp. cells and corresponding (poor) segmented image. B) Denoised image (BM3D, sigma = 15) and corresponding improved cells masks covering the visual cell shapes and sizes.

### Isolated Dimalis work flow

Finally, the Dimalis pipeline was wrapped in a *docker* container to ‘freeze’ the version of each underlying package. This presents the strong advantage that launching Dimalis on the same time-lapse images with the same set of parameters will always lead to the same results, regardless of updates of the packages that it relies on. To illustrate this, we launched four different versions of the Omnipose cell segmentation tool on a common set of images (Supplementary figure 1). The results clearly demonstrated that segmentation is drastically influenced by the version of the underlying cell segmentation (the version 0.7.1 being maintained in the Dimalis package).

### Extracting cell growth parameters from Dimalis

Potential applications of Dimalis lie in the extraction of cell growth parameters from time-resolved imaging data. We followed individual microcolony growth of four random soil isolates (*Burkholderia, Curtobacterium, Mucigalinibacter* and *Rahnella* spp., Fig. 5A) from single seeded cells in monoculture on agarose patches with the same added nutrients, by time-lapse imaging with 10-minute intervals. Both the faster (e.g., *Rahnella*) and slower-growing strains (i.e., *Mucigalinibacter*) follow Monod-type biomass growth (here taken from the increase of the natural logarithm of the sum of the areas of the cells belonging to a lineage over time, Fig. 5B), and logarithmic population growth (deduced from the increase in number of segmented cells per lineage over time, Fig. 5C). This allows for detailed measurements of maximum exponential growth rates from individual lineages based on cell area (roughly equivalent to biomass, *μ*_max_; Fig. 5D), or maximum exponential division rates (taken from the log_2_ of the population size increase; *k*_max_, as in Fig. 5E). Indeed, the mean *μ*_max_ is roughly equivalent to ln(2) × *k*_max_ as growth kinetic theory would stipulate. When broken down to cell division times (the time differences between subsequent divisions per generation), one can see that there is quite some individual variation of cell division within lineages (Fig. 5F). Note that on average, the first generation time is slightly shorter, which is likely due to the time needed to mount the chamber with the surface-grown cells under the microscope and start the imaging process. Two of the four strains, *Curtobacterium* and *Rahnella*, show relatively constant cell division times, whereas lineages of *Burkholderia* and *Mucigalinibacter* have a tendency of longer generation times for the first 2–5 cell divisions than at later time points in the lineage (Fig. 5F). This example illustrates, therefore, the type of extractable data from time-lapse microcolony imaging to study differences in individual cell behaviour.

**Figure 5:**
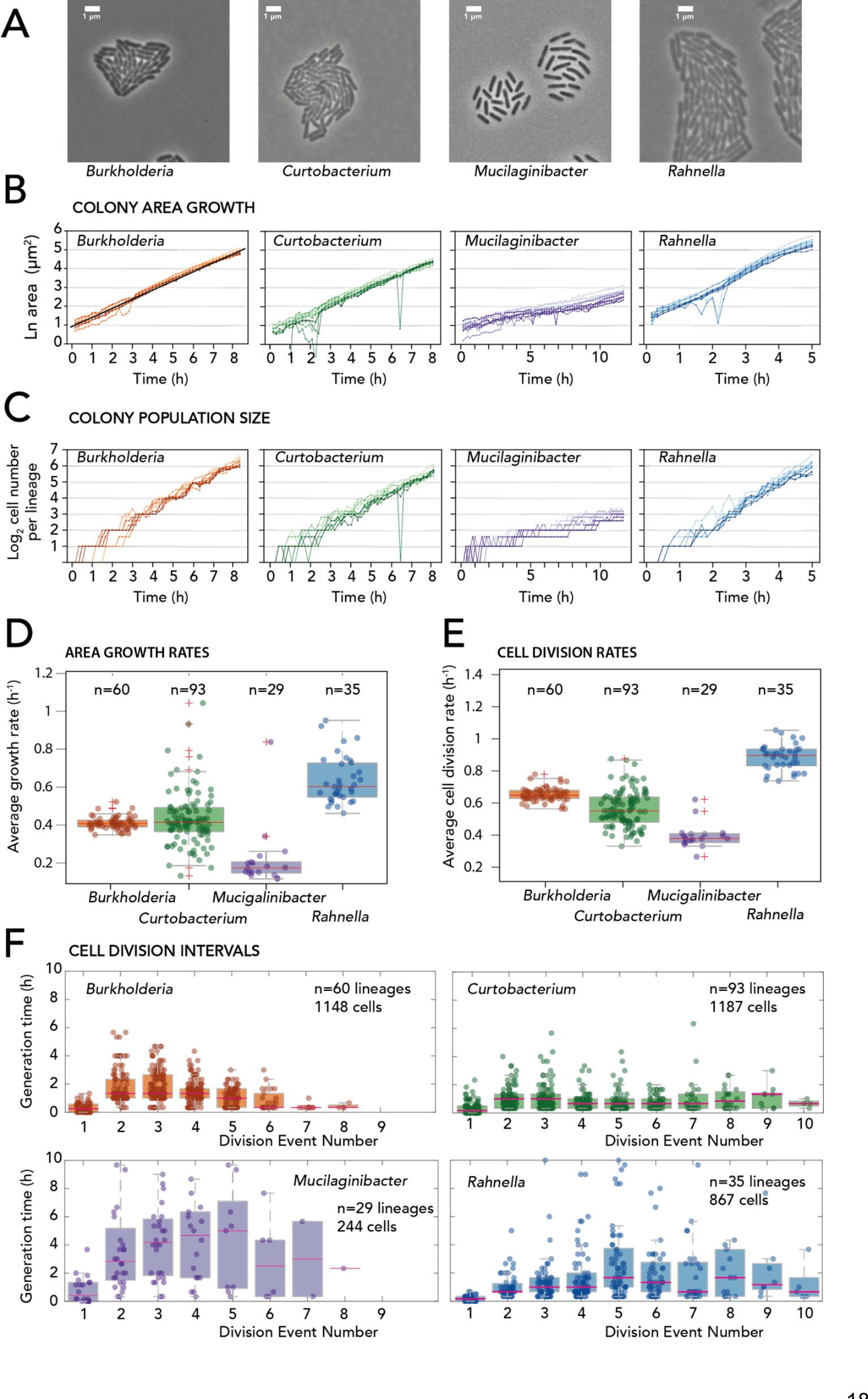
Growth kinetics of *Burkholderia, Curtobacterium, Mucigalinibacter* and *Rahnella* from microcolony time-lapse imaging. A) Micrographs of microcolonies of each of the four bacterial strains growing on complex medium after ca. 6 – 8 h (bar represents 1 μm). B) Natural log of the sum of the segmented cell areas in a genealogy lineage over time for the four different strains. Lines indicate 10 individual lineages. Note the difference in time scale for *Mucigalinibacter* and *Rahnella* compared to *Burkholderia* and *Curtobacterium*. C) Same as (B) but now for the lineage sizes as cell numbers, displayed as the log_2_ of the sum of cells per lineage over time. Individual lines and colour shades represent the 10 different microcolony lineages. D) Average lineage expansion rates, calculated as the mean of slopes across a moving interval of 15-20 time points, fitted to the natural logarithm of the lineage area increase over time (as in B), taking only those slopes with an R^2^ higher than 0.89 into account. Boxplots show the 25^th^, median, and 75^th^ percentiles. n = number of individual lineages (shown as individual dots). E) As (D), but with the average cell division rates, calculated from the lineage cell population size increase (slopes from C). Note how roughly the mean expansion rates correspond to ln(2)×the cell division rates, according to theory. F) Generation times for each of the four strains over the first ten observed divisions. Boxplots are overlaid with the values from cells in individual lineages (as dots). Note how the first generation time is shorter because of the time to place the cell holder under the microscope and start imaging.

### Quantifying cell division rates in microfluidics chambers

To illustrate further applications of Dimalis on biological questions, we controlled bacterial growth in microfluidics chambers under constant nutrient flow and aimed to extract cell division rates as a function of spatial position in the chamber. Bacterial cell growth in microfluidics typically results in very crowded images, thus requiring efficient cell segmentation tools to detect cell contours in an automated way and over time. Figure 6A shows segmentation and tracking of *Pseudomonas* sp. Leaf 15 growing on 10 mM xylan at 0.1 ml h^−1^ flow rate in a microfluidic device. Tracking indicates the directionality of cell divisions as a function of flow and cell elongation rates (in h^−1^), which are retrieved by exponential fitting of the increase of the long cell axis over time (images every 10 min). The spatial elongation rate is then displayed as the sum per 15×15 μm^2^ tiles (Fig. 6B) and averaged across columns. In this setting, cells divide homogenously in space (Fig. 6C), indicating that their growth was not influenced by any gradient of nutrients or oxygen along the chamber. When imposing a nutrient gradient on the microfluidic chamber, one can see how the averaged elongation rates increase close to the gradient source and decrease as a function of distance (Fig. 6D and E).

**Figure 6:**
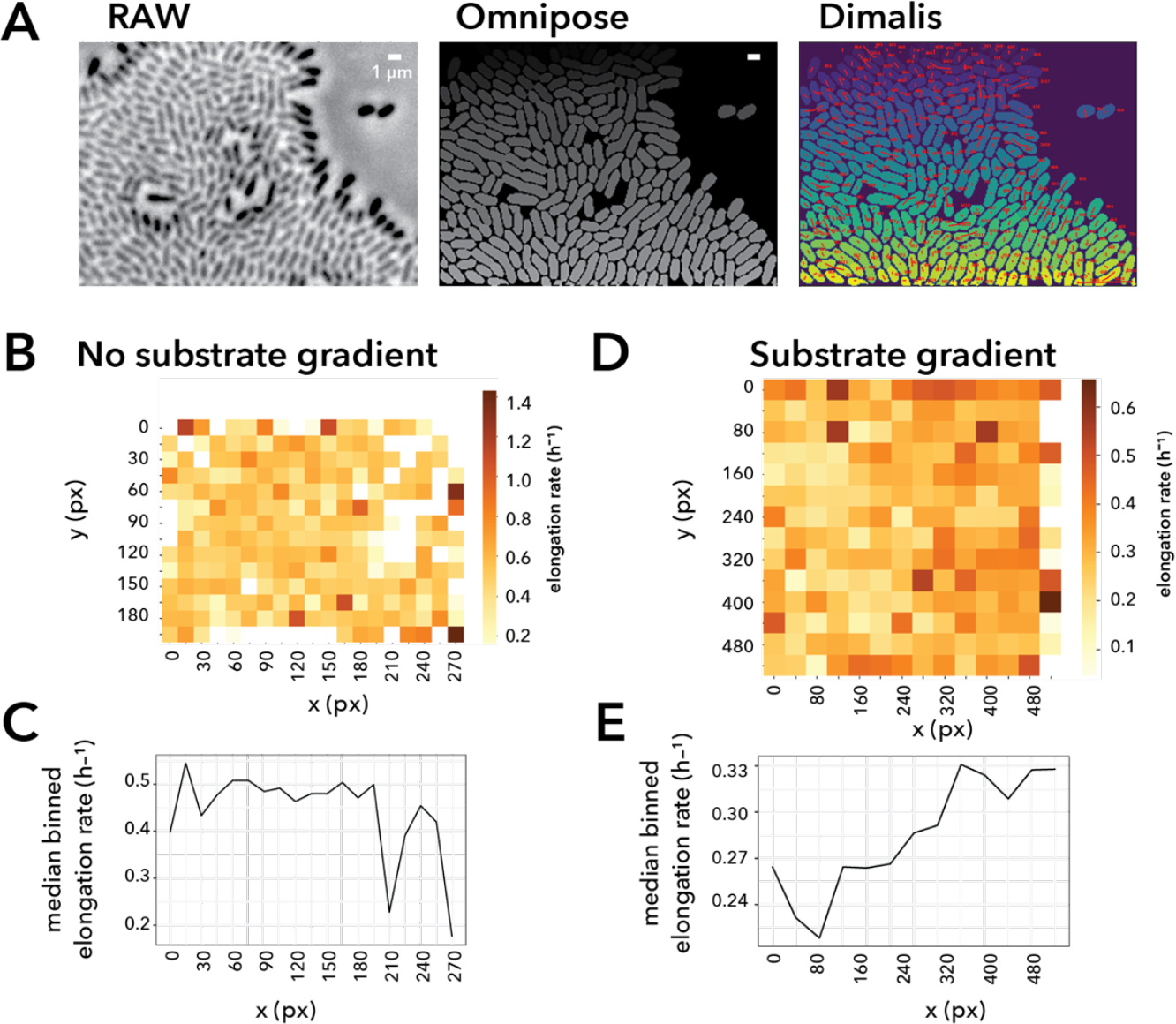
Cell elongation rates in microfluidics as function of substrate . A) Phase-contrast image of quasi-discontinuous monolayer growth of *Pseudomonas* sp. Leaf15 in a microfluidics chamber with corresponding cells masks resulting from the cell segmentation in Dimalis. Note how cell tracks leading from the previous to the current image are overlaid on the image with a small red line. B) Measured elongation rates over the image, summarized per 15×15 pixel bins, here in absence of substrate gradient imposed on the cells. C) Median binned elongation rates as a function of distance (in pixels) to the left image boundary. D) and E), as B but in presence of a xylan substrate gradient (from left to right).

### Extracting cell lineages displaying bistable switching to transfer competent cell subpopulations

In the third example, we followed growth and subsequent stationary phase of individual *P. putida* UWC1-ICE*clc* cells on agarose disks (1 mm thickness) with 3-chlorobenzoate as sole carbon and energy source to detect formation of transfer competent cells. The integrative and conjugative element ICE*clc* is stochastically activated in a subpopulation of cells in stationary phase (Sulser, et al., 2022), which can be detected by expression of a single copy chromosomally-integrated transcriptional reporter fusion between the promoter of the *inrR* gene of the ICE and *mCherry* (Sulser, et al., 2022). Only cells expressing *inrR* become transfer competent for ICE*clc* conjugation. In addition, the strain carries a single-copy reporter between the promoter of the *ahpC* alkylhydroxyperoxidase gene and *egfp* as a monitor for the appearance and levels of oxidative stress. Dividing cells were imaged every 30 min for up to 96 h in phase-contrast, and with additional mCherry and GFP fluorescence imaging. In the example shown here, one can see how a small number of cells with very bright mCherry fluorescence are visible within microcolonies imaged at t = 72 h (representative for the transfer competence status, Fig. 7A). Tracing individual cells over time showed that the two brightest cells appear around 25 h after incubation start, and that four further individual cells appeared with intermediate mCherry levels (Fig. 7B). No direct correlation of individual cell mCherry and GFP fluorescence was observed (Fig. 7B). Cell lineage plotting traces the appearance of transfer competent cells (based on their mCherry signal) to early stationary phase, when cells have stopped dividing. In contrast, oxidative stress signals are highest at the earliest time points of cell division, and with a second smaller increase at the onset of stationary phase (Fig. 7C). This example thus shows how cell lineage data can be extracted to quantify subpopulation events.

**Figure 7:**
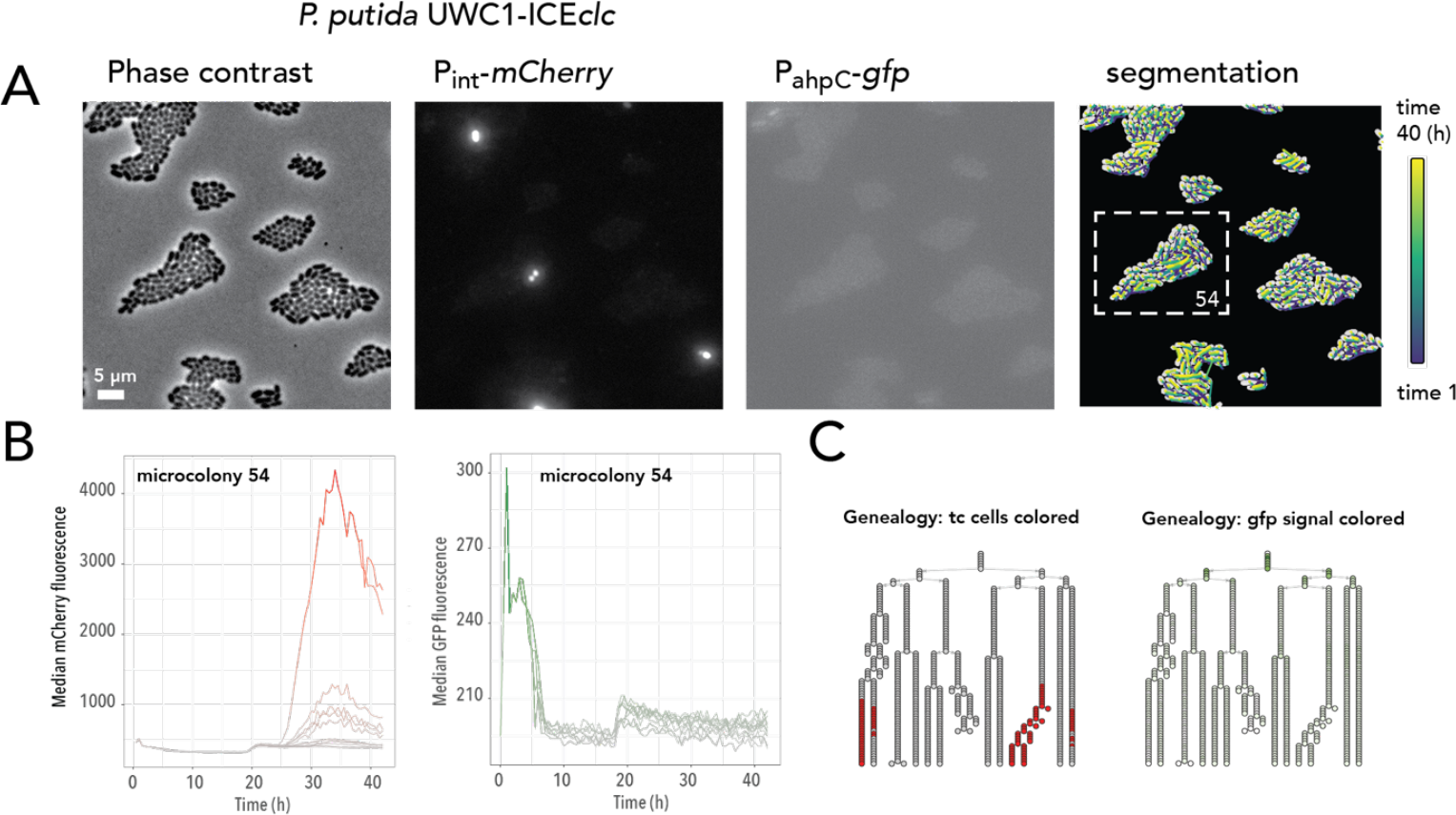
Double fluorescence reporter analysis in *P. putida* microcolonies. A) Microscope images of *P. putida* microcolonies grown on 3-chlorobenzoate to stationary phase (32 h), in phase contrast, mCherry fluorescence (from a single copy transcriptional fusion to the P_int_-promoter of the integrative and conjugative element ICE*clc*), GFP fluorescence (from a single copy transcriptional fusion to the *ahpC* oxidative stress promoter), and a segmentation overlap of the grown colony at all time points. B) Detail of mCherry and GFP fluorescence development over time of microcolony 54 (boxed in A), indicating few brightly appearing transfer competence cells in stationary phase, and initially high oxidative stress response in dividing cells. C) Derived microcolony genealogy with dividing cells, each cell and time point being represented by a small circle. Colors indicate median fluorescence values above the 95th percentile. Note how genealogies can be confused by small tracking differences in stationary phase colonies, resulting in a horizontal repositioning of circles.

## DISCUSSION

We present and tested here an automated pipeline for segmentation and tracking of bacterial cell types, to facilitate analysis of time-lapse microscopy imaging and extraction of relevant growth kinetic data. The pipeline was combined mostly from existing individual software tools, notably BM3D (Djurović, 2016; Sheng, et al., 2014), Omnipose (Cutler, et al., 2022) and STrack (Todorov, et al., 2023), wrapped inside a Docker structure for consistency and ease of application. We minimize user input, restricting it to a single parameter setting for the denoising, and three parameters that cover the expected average cell diameter, the expected distance between cells for the tracking and the allowed division angle. The pipeline organizes the output data and produces a straightforward summary .csv output table that can serve as input for subsequent analysis steps. As we demonstrated here, the pipeline extracts useful data from a number of different bacterial cell types based on phase-contrast imaging, and in different settings (e.g., microfluidics, microcolony growth). It can also extract corresponding fluorescent imaging data, useful for, e.g., reporter gene expression studies. The tools work well even on relatively crowded cell-images, although we do find that segmentation and tracking become increasingly imprecise with e.g., stationary phase microcolonies with > 200 cells. We also experienced that occasional blurred images, arising from long time series with automated focusing, can only be partly resolved by denoising. Therefore, poor quality images will still lead to poor quality results. We acknowledge that a very recent new segmentation and tracking program was presented, which may resolve recurrent issues in positioning of time-lapse images, but which we could so far not test or implement ourselves (Ollion, et al., 2024). Larger image position shifts in time-series remain problematic for the cell tracking, in which case repositioning packages such as pyStackReg (Thevenaz, et al., 1998) may be necessary to be included on the image stacks before feeding them into Dimalis.

We welcome feedback on Dimalis usage and hope the tools can facilitate entry users to start with single cell bacterial image analysis.

## ACKNOWLEDGEMENTS

This work was supported by the Swiss National Science Foundation (Sinergia program, grant CRSII5 189919/1 to J.M.), and by the National Centre in Competence Research (NCCR) in Microbiomes (grant number 180575 to J.M.). ARP is supported by a James. S. McDonnell Postdoctoral Fellowship (2020-1332). The funders had no role in study design, data collection and interpretation, or the decision to submit the work for publication.

