## Supplementary figure 1 for "Dimalis: A complete standalone pipeline to analyse prokaryotic cell growth from time-lapse imaging"

**Supplementary Figure 1: Effect of different program versions of cell segmentation**. A) Phase-contrast images corresponding to four successive time points of a growing *P. putida* microcolony. B) Obtained segmentation masks with four different versions of Omnipose (0.7.0, 0.7.1, 0.7.2 and 2.2) on the images in (A) with the same trained model and the same parameters.

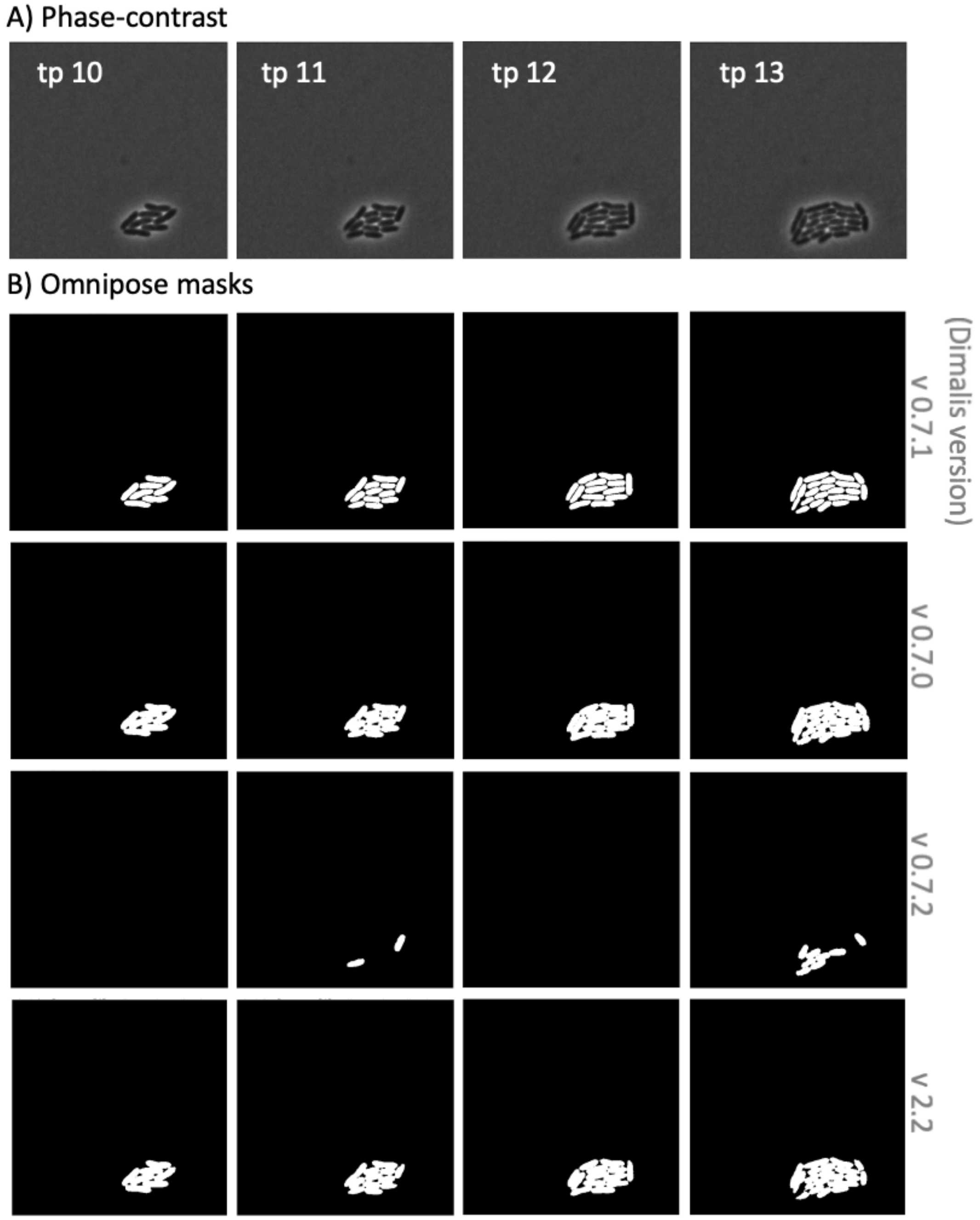
